# Polymeric nanoparticles delivery of AMPK activator 991 prevents its toxicity and improves muscle homeostasis in Duchenne Muscular Dystrophy

**DOI:** 10.1101/2024.01.16.575840

**Authors:** Ilaria Andreana, Anita Kneppers, Sabrina Ben Larbi, Federica Tifni, Aurélie Fessard, Jaqueline Sidi-Boumedine, David Kryza, Barbara Stella, Silvia Arpicco, Claire Bordes, Yves Chevalier, Bénédicte Chazaud, Rémi Mounier, Giovanna Lollo, Gaëtan Juban

## Abstract

Muscular dystrophies, such as Duchenne muscular dystrophy (DMD), are caused by permanent muscle injuries leading to chronic inflammation. In that context, macrophages harbor an altered inflammatory profile that contributes to fibrosis through the secretion of the profibrotic cytokine TGFβ1. We previously showed that AMP-activated protein kinase (AMPK) activation reduces TGFβ1 secretion by macrophages and improves muscle homeostasis and muscle force in a mouse model of DMD. This makes AMPK an attractive therapeutic target for treating chronic inflammation and fibrosis in DMD. However, potent direct AMPK activators like compound 991 show strong adverse effects *in vivo,* preventing their direct use. Here, we encapsulated 991 into biodegradable polymeric poly(lactic-*co*-glycolic) acid (PLGA) nanoparticles for *in vivo* delivery, in an attempt to overcome toxicity issues. We show that 991-loaded PLGA nanoparticles retained drug activity on fibrotic macrophages *in vitro*, by reducing their secretion of TGFβ1. In the D2-mdx pre-clinical DMD mouse model, intravenously injected PLGA nanoparticles reached gastrocnemius and diaphragm muscles, which are the most affected muscles in this model. Chronic intravenous injections of 991-loaded PLGA nanoparticles decreased inflammation in both muscles, which was associated with fibrosis reduction and increase in myofiber size and muscle mass in the gastrocnemius. No impact on blood cell counts and liver enzymes was observed. These results demonstrate that nanomedicine is an efficient strategy to deliver AMPK activators *in vivo* to target inflammation and improve the dystrophic muscle phenotype.

## Main

Duchenne Muscular Dystrophy (DMD) is a severe and progressive muscle-wasting disease leading to loss of ambulation, need for mechanical respiratory assistance and premature death at around age 30 years^1,2^. Mutations observed in DMD induce a lack of the Dystrophin muscle isoform^1,2^, that belongs to the dystroglycan complex stabilizing the link between the cytoskeleton and^3^. This leads to myofiber fragility and causes permanent muscle injuries and attempts of regeneration, chronic inflammation and fibrosis. Fibrosis is a critical component of the disease as there is a strong correlation between the level of muscle fibrosis and the loss of muscle function^4^. Currently, there are no curative treatments for DMD and most of the research is directed towards the development of gene therapy strategies to rescue the expression of Dystrophin in patients^5^. Standard palliative care for patients consists in the chronic administration of glucocorticoids. Although they have been shown to delay disease progression and ambulation loss^6^, long term exposure to glucocorticoids is associated with strong adverse effects including obesity, metabolic dysregulation and myofiber atrophy^7,8^.

The chronic inflammation observed in DMD is characterized by the presence of pro-inflammatory and pro-fibrotic macrophages that differ from those observed after an acute damage. Attempts to inhibit monocyte entry into the DMD muscle was shown beneficial for a couple of weeks^9,10^, but deleterious at longer term^11^, indicating that macrophages are needed for muscle repair in DMD, as they are after an acute damage^12^. In normal skeletal muscle, the damage triggers blood monocyte entry into the injured area where they become pro-inflammatory macrophages favor muscle stem cell (MuSC) proliferation^13,14^ and limit fibro/adipogenic progenitor (FAP) expansion^15,16^. The resolution of inflammation is triggered by their switch towards an anti-inflammatory/restorative profile^13^ which stimulates MuSC differentiation and fusion^14^, favors extracellular matrix (ECM) production by fibroblastic cells^15,16^ and promotes angiogenesis^17^. We identified that activation of the metabolic sensor AMPK is required for the macrophage inflammatory switch^18–20^. In DMD, the pro-inflammatory macrophages are associated with fibrosis while anti-inflammatory macrophages are associated with myogenesis^16,21^. In this context, pharmacological skewing of pro-inflammatory/pro-fibrotic macrophages towards a pro-resolutive phenotype using indirect activators of AMPK such as Metformin^16^ or NaHS^22^ reduces muscle fibrosis and improves dystrophic muscle homeostasis and function *in vivo* in a DMD mouse model, identifying AMPK as a potential therapeutic target.

In the perspective of targeting AMPK, the use of a direct activator appears to be more relevant as indirect activators may also activate other effectors, potentially leading to uncontrolled adverse effects. However, with the notable exception of the PXL770 compound^23^, direct AMPK activators performed poorly in clinical trials either because of a lack of patient condition improvement or because of adverse effects^24,25^. Among them, the synthetic compound 991 (also known as Ex229) is a cyclic benzamidole derivative identified as a potent allosteric activator of AMPK in skeletal muscle *ex vivo*^26^ that inhibits TGFβ secretion by fibrotic macrophages *in vitro*^16^. Therefore, it represents a good candidate to selectively and potently activate AMPK in DMD muscle to modulate inflammation, dampen fibrosis and improve muscle function. While the literature does not report any study describing the use of 991 *in vivo*, it is accepted that 991 injection in mice induces strong adverse effects such as hemolysis, platelet aggregation and liver alteration (unpublished observations), preventing its direct use.

Recently, gold^27^ or mesoporous silica^28^ nanoparticles (NPs) have been used to target muscles in dystrophic mouse models. However, these type of NPs can be associated with toxic effects^29,30^. To enable the AMPK activator 991 delivery *in vivo,* we optimized its encapsulation into biodegradable polymeric poly(lactic-*co*-glycolic) acid (PLGA) nanoparticles. PLGA nanoparticles exhibit a small size (<100 nm) that allows them to enter passively and accumulate into inflamed tissues through capillaries thanks to the enhanced permeation and retention (EPR) effect^31^. Moreover, they are FDA and EMA approved for intravenous administration and have already been used in clinical trials for the delivery of anti-cancer treatments^32^. In the context of DMD, this administration route appears to be the most relevant as it potentially allows to target all the affected muscles throughout the body.

Here, we investigated the possibility to deliver the AMPK activator 991 *in vivo* through its encapsulation into PLGA NPs in the D2-mdx preclinical mouse model, which exhibits a severe dystrophy^33–35^. We first assessed the feasibility of encapsulating 991 into PLGA NPs via a continuous scalable process, namely microfluidics, to obtain a reproducible and highly efficient incorporation of the drug in the NPs while guaranteeing the automation of NP formation. Moreover, we produced a dry form of the loaded-PLGA NPs to increase their storage stability over 21 days and reach a complete elimination of residual organic solvents. We next showed that 991 retained its biological activity towards fibrotic macrophages *in vitro* when loaded into PLGA NPs. Then, we demonstrated that upon intravenous injection, PLGA NPs reached both gastrocnemius and diaphragm muscles of the D2-mdx mouse where they are internalized by muscle cells. Finally, chronic intravenous injections of 991-loaded PLGA NPs decreased the number of macrophages in both muscles, which was associated with improved muscle homeostasis in the gastrocnemius.

## Results

### Optimization of 991-loaded PLGA nanoparticles by continuous formulation process using DoE approach

NPs were successfully formulated by both conventional manufacturing method (nanoprecipitation) and microfluidic technique (Table S1). Based on the data obtained by nanoprecipitation (NPs in the range of 150–170 nm, polydispersity index (PdI) in the range of 0.05–0.18 and high encapsulation efficiency (EE) of 85%), we produced PLGA NPs using a microfluidic system to benefit from controlled flow condition and the efficient mixing at high flow rates of toroidal micromixer (Fig.1A). Polymer nanocarriers were prepared by setting a total flow rate (TFR) of 12 mL/min and a flow rate ratio (FRR) of 1:2 to obtain sub-100 nm NPs (Fig.1B). In the controlled-mixing process, NPs were characterized by a size of 70 and 81 nm, for empty and loaded nanosystems, respectively, with a slight diameter change in the presence of the encapsulated drug. Furthermore, Transmission electron microscopy (TEM) confirmed the sub-100 nm mean diameter and highlighted the rounded shape of microfluidic-prepared NPs (Fig.1C). PLGA NPs are widely described in the literature for their ability to encapsulate various drug molecules^36,37^. However, the impact of FRR, polymer and drug concentration, in this case 991 concentration, were investigated for the microfluidic optimization to define a roadmap for a continuous production of NPs via design of experiment (DoE) approach. EE and drug loading (DL) were identified as critical quality attributes that are related to quality product. Using a TFR of 12 mL/min, we obtained NPs characterized by a mean diameter below 150 nm and a PdI around 0.2 (Table S2), as shown by previous results of polymer NP preparation at high TFR (10–15 mL/min)^38,39^. The fitting quality of the developed models for EE and DL was confirmed by the determination coefficients R^2^ and the residual error value of each experiment, within the range of 2xSD^exp^ (Table S3). The results that we obtained from the DoE evidenced the influence of operative and formulative parameters on the size, EE and DL loaded PLGA NPs. Regarding the EE (Fig.1D), the DoE results showed that it is maximized at low polymer concentration (5 mg/mL) since the amount of polymer is not enough to entrap the drug in a high amount. As expected, DL increased with the drug concentration and by decreasing the polymer amount. DL was slightly more influenced by the drug amount than the polymer concentration (Fig.1E).

**Figure 1.**
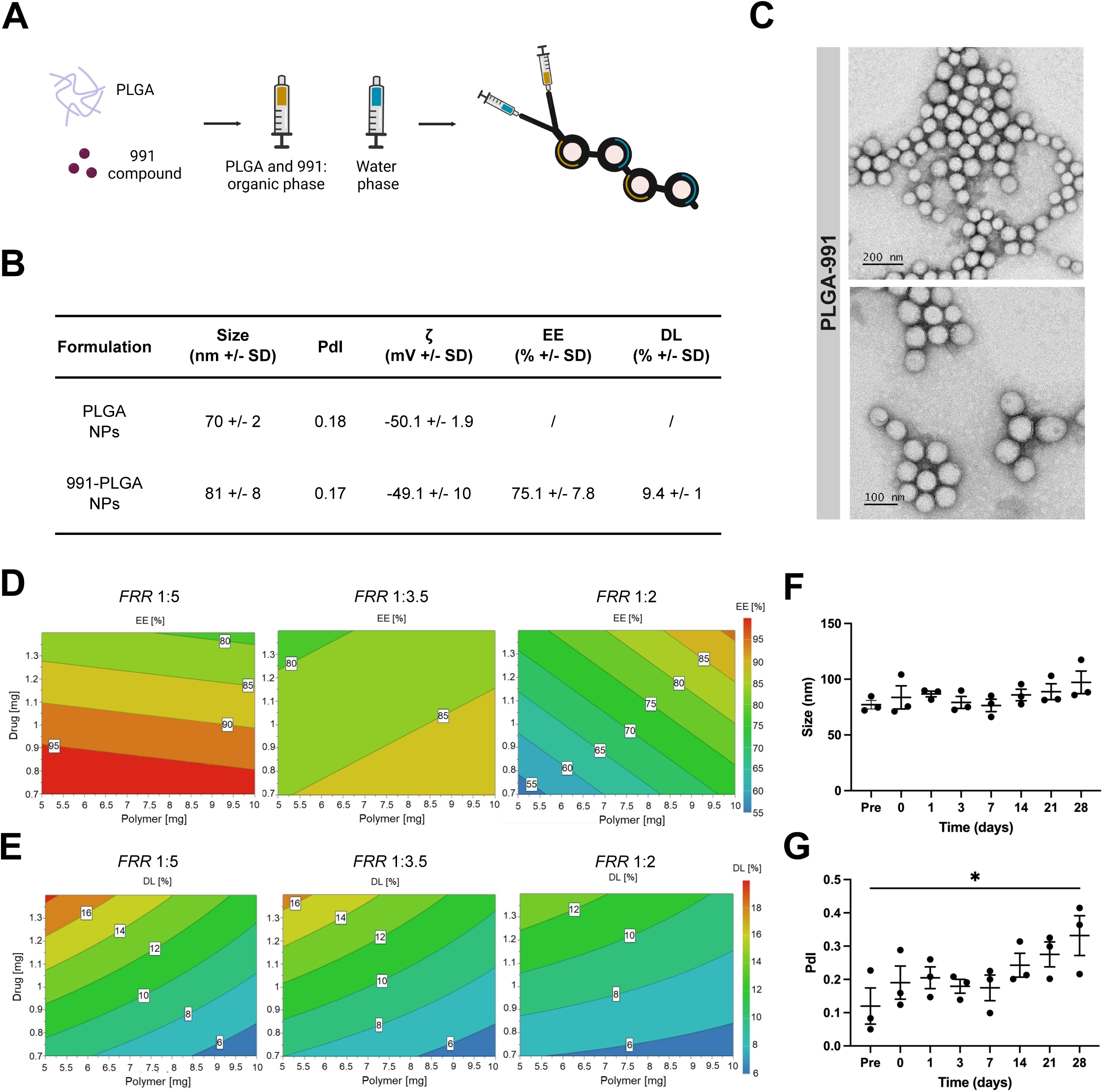
Manufacturing translation of 991-loaded PLGA NPs: formulative optimization of a continuous production process. (A) Schematic representation of the microfluidic micromixing. (B) Polymer NP physicochemical analysis. Values are given as mean ± SD (n = 3). (C) TTEM of empty PLGA NPs (top) or 991-loaded PLGA NPs (bottom). (D-E) Contour plots of the response surface for EE (D) and DL (E) at FRR 1:5 (left panels), 1:3.5 (middle panels) and 1:2 (right panels). (F-G) Characterization of 991-loaded NPs after freeze-drying. Analysis of the size (F) and PdI (G) upon resuspension of loaded freeze-dried NPs.

The best compromise in terms of EE and DL was obtained by working between FRR 1:2 and 1:3.5 (experiments 8, 9 and 10) (Table S2). Thus, keeping satisfying results for EE and DL, we decided to investigate the biological activity of the formulation prepared at FRR 1:2 and with 1 mg/mL of 991 and a low PLGA concentration of 6 mg/mL that allows reducing NP mean diameter.

For deeper consideration about the pharmaceutical application of sub-100 nm loaded-PLGA NPs, NMR analysis was performed to evaluate the presence of residual organic solvents which could impact the stability and biocompatibility of PLGA formulation. 1H NMR spectra analysis showed residual of acetonitrile even after NP purification after dialysis (data not shown). For this reason and to obtain a dry form of NPs improving their storage stability, the formulation was freeze-dried after solvent elimination in the presence of 1 to 12 % (w/v) of trehalose as preferential cryoprotecting agent^40^. After freeze-drying, all the loaded formulations had a non-collapsed cake aspect, allowing an easy resuspension in Milli-Q water. In addition, the 12% w/v of trehalose allowed NPs to retain the size and shape they had before freeze-drying for up to 21 days (Fig.1F-G, Fig.S1), and a suitable osmolarity (286 mOsm) for *in vivo* administration (Table S4).

### PLGA encapsulation of the 991 AMPK activator retains its anti-fibrotic properties

In order to evaluate the impact of NP encapsulation on 991 drug activity, we first tested the polymer nanosystem *in vitro* on bone marrow-derived macrophages (BMDMs) that were activated into fibrotic macrophages by fibrotic muscle protein lysates^16^ (Fig.2A). First, BMDM incubation with PLGA NPs loaded with DiD fluorescent dye showed their fast internalization by macrophages, as soon as 2 h and that lasted up to 24 h (Fig.2B). The activity of encapsulated 991 was then assessed by monitoring the phosphorylation of the two AMPK targets acetyl-CoA carboxylase (ACC) and regulatory associated protein of MTOR (RAPTOR). As expected, BMDM treatment with 10 or 20 µM free 991 increased the phosphorylation of ACC (+185 and +230% *vs*. Vehicle-treated control, respectively) and RAPTOR (+208 and +329% *vs*. Vehicle-treated control, respectively) (Fig.2C-D). Notably, BMDMs treated with 991-loaded PLGA NPs (PLGA-991) at 10 and 20 µM showed a similar increase in the phosphorylation of ACC (+192 and +146% *vs*. PLGA-treated control, respectively) and RAPTOR (+128 and +96% *vs*. PLGA-treated control, respectively) (Fig.2C-D), indicative of AMPK activation. The capacity of PLGA-991 to inhibit TGFβ1 production by fibrotic BMDMs was then quantified by ELISA. As we previously showed^16^, 991 treatment inhibited the secretion of TGFβ1 by fibrotic BMDMs (−65% *vs*. Vehicle-treated control) (Fig.2E). A similar reduction of TGFβ1 production was observed following incubation with PLGA-991, -59% *vs*. PLGA-treated control) (Fig.2E). Importantly, the decrease was not due to a potential toxic effect, as PLGA-991 treatment even increased cell viability (Fig.S2).

**Figure 2.**
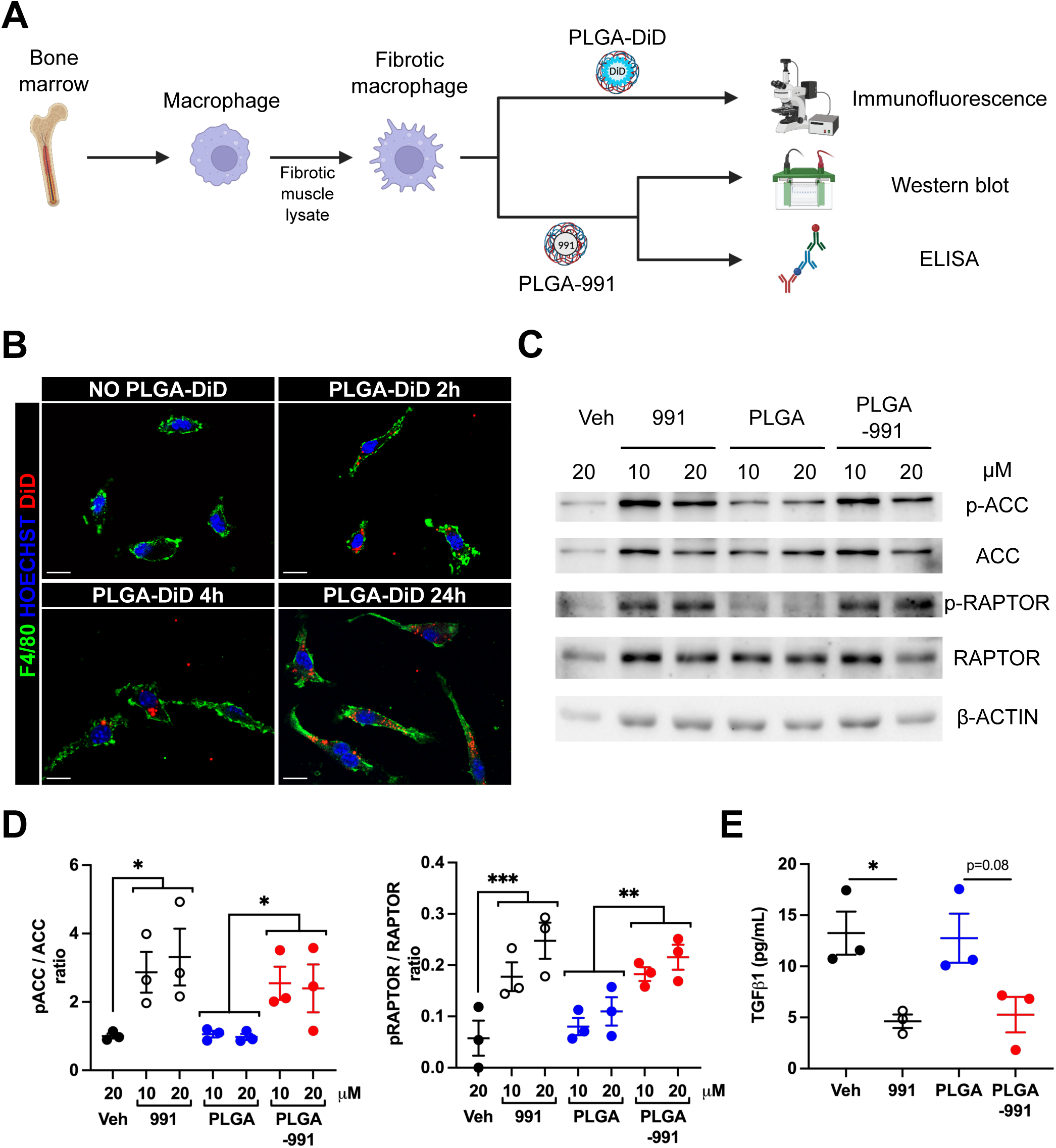
PLGA encapsulation of 991 AMPK activator retains its anti-fibrotic properties. (A) BMDMs were activated into fibrotic macrophages and incubated with various types of PLGA NPs. (B) Cells were incubated without or with DiD-loaded NPs (PLGA-DiD) for 2, 4 and 24 h, immunolabeled with anti-F4/80 antibody and nuclei stained with Hoechst. A representative image of each time point is shown. Scale bar: 10 μm. (C-E) Fibrotic macrophages were treated without (Veh) or with 10 or 20 µM of 991 alone, empty PLGA or PLGA-991 for 6 h and the phosphorylation of ACC and RAPTOR was quantified by immunoblot. (C) Representative immunoblots. (D) Quantification of the ratio between the phosphorylated over the total ACC (left) or RAPTOR (right) proteins. (E) Fibrotic macrophages were treated without (Veh) or with 10 µM of 991 alone, empty PLGA or PLGA-991 for 20 h and the TGFβ1 production was measured by ELISA.

These results show that 991 encapsulation into PLGA NPs allows its internalization by fibrotic macrophages and retains its full activity.

### PLGA nanoparticles efficiently target diaphragm and gastrocnemius muscles in D2-mdx mice

In order to assess whether small-size PLGA NPs (≤100 nm) can reach and accumulate into skeletal muscles, fluorescent PLGA-DiD NPs were administered by intravenous injection into D2-mdx mice and their biodistribution was assessed by imaging (Fig.3A). As previously described in other mouse models^41^, we observed a rapid accumulation of PLGA NPs in the liver, spleen and lungs (Fig.3B-C). In the liver, NP residence lasted over 72 h, but the fluorescence intensity decreased over time in the spleen (17% of initial fluorescence remaining after 48 h) and lungs(28% of initial fluorescence remaining after 24 h) (Fig.3C), indicative of an elimination of the nanosystem. Interestingly, fluorescence was also detected in thè diaphragm as soon as 0.5 h, although at a lower level as compared with the other organs, with an almost complete clearance after 24 h (Fig.3B,D). Flow cytometry analysis showed that the fluorescence in the diaphragm was associated with NP uptake by the cells of the muscle (9.6% and 10.3% of the total muscle cells at 0.5 and 1 h post-injection, respectively (Fig.S3 and Fig.3E). The nanosystem signal in the cells decreased over time (3.7% of cells after 72h) (Fig.3E), indicative of its metabolization by the cells. Finally, we investigated the gastrocnemius, which is another muscle strongly affected in the D2-mdx model. Even though the PLGA-DiD fluorescence was not detected on the whole gastrocnemius muscle by imaging (not shown), we observed NP uptake by the cells (7% of the muscle cells at 0.5 h post-injection), with a reduction after 48 h (3.7% of cells) ( Fig.3F).

**Figure 3.**
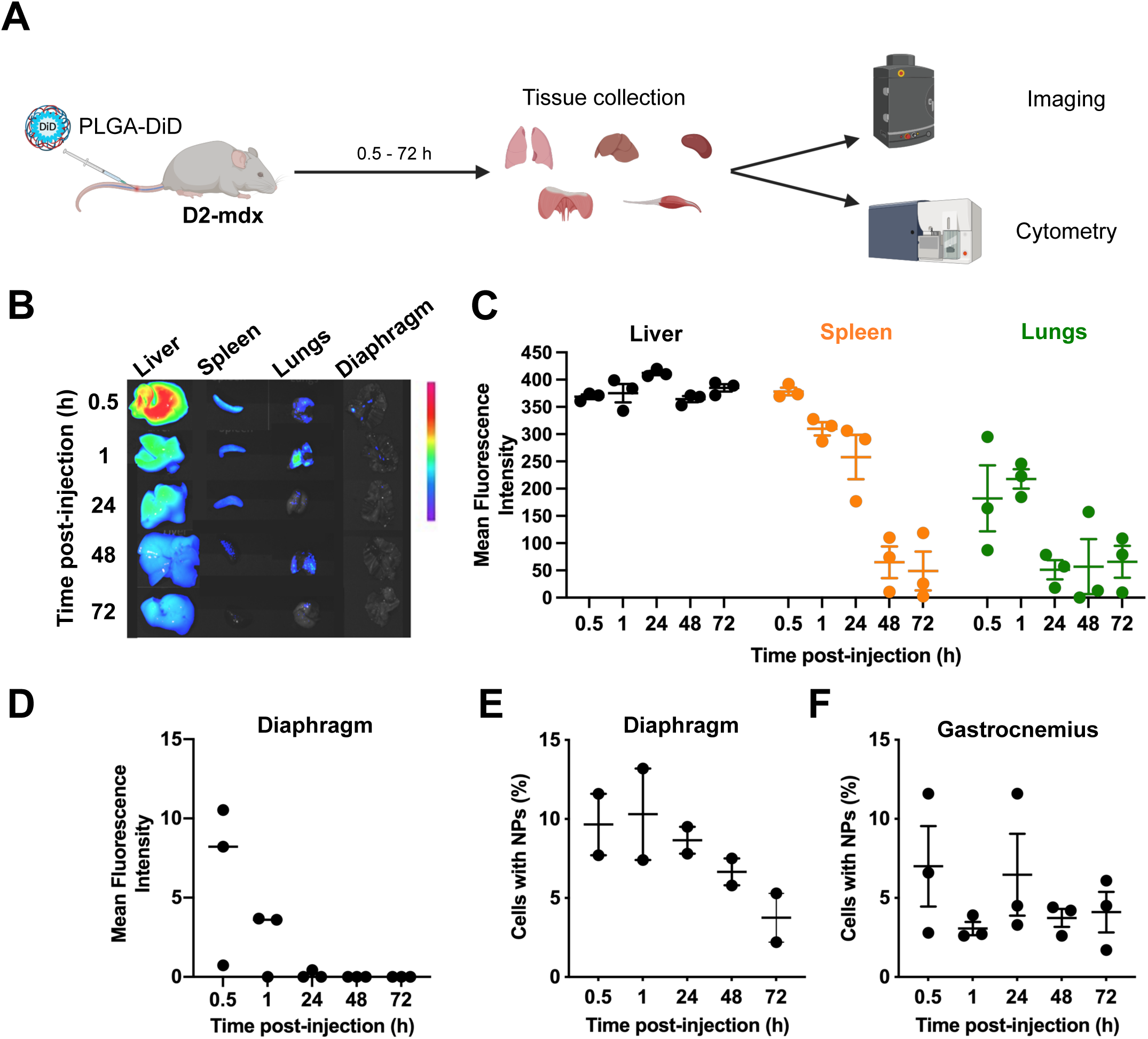
Biodistribution of PLGA NPs in D2-mdx mice. (A) D2-mdx mice were injected intravenously with DiD-loaded PLGA NPs and sacrificed 0.5, 1, 24, 48 and 72h later. (B-D) Organs were harvested and imaged to quantify DiD fluorescence. (B) Representative images of imaged organs. (C) Mean fluorescence intensity quantified in liver (black), spleen (orange), and lungs (green). (D) Mean fluorescence intensity quantified in the diaphragm muscle. (E-F) Diaphragm (E) and gastrocnemius (F) muscles were dissociated and the percentage of cells having internalized DiD-loaded PLGA NPs was determined by flow cytometry.

These results show that PLGA NPs efficiently reach the diaphragm and gastrocnemius muscles of D2-mdx mice where they are internalized by the cells.

### Chronic PLGA-991 treatment does not induce adverse effects

In order to evaluate the potential *in vivo* toxicity, PLGA-991 NPs were administrated by chronic (every other day) intravenous injections over a time period of 21 days, then blood, liver, spleen and lungs were harvested (Fig.4A). We did not observe any effect of the PLGA-991 treatment on liver, spleen and lung mass as compared with PBS or empty PLGA NPs (Fig.4B). Similarly, PLGA-991 chronic administration was not associated with a modification of the number of erythrocytes, platelets and leukocytes in the blood (Fig.4C), indicative of an absence of blood toxicity. Finally, we monitored potential liver alteration by quantifying the plasma level of alanine (ALT) and aspartate (AST) transaminases, which are two enzymes that are released in the blood upon liver injury. Importantly, neither of these enzymes were increased in the plasma of animals treated with PLGA-991 NPs as compared with PBS or empty PLGA NPs (Fig.4D,E).

**Figure 4.**
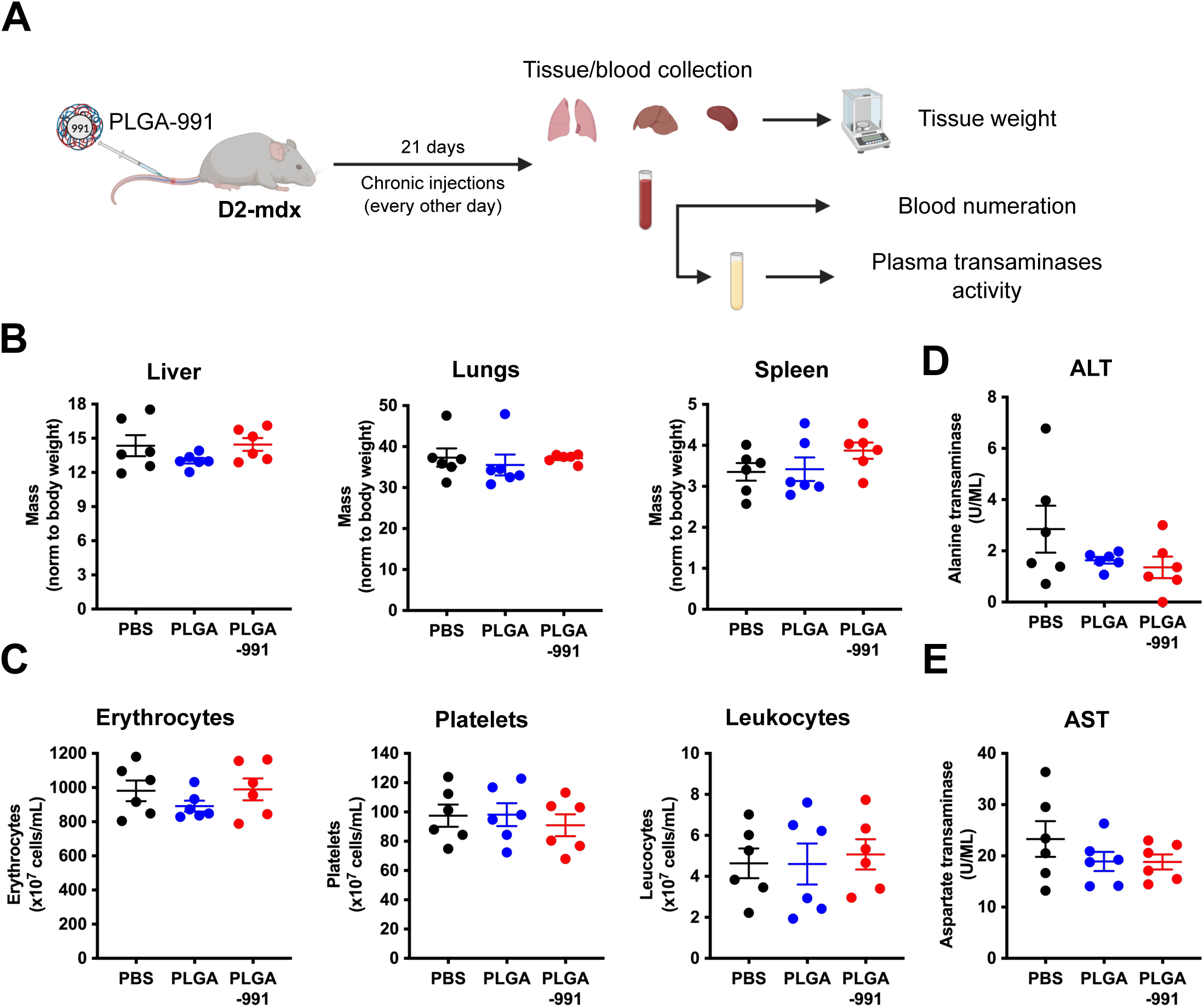
Chronic PLGA-991 treatment does not induce adverse effects. (A) D2-mdx mice were treated with chronic intravenous injections of PBS, PLGA NPs or PLGA-991 for 21 days. Liver, lungs and spleen were harvested and blood collected. (B) Relative mass of liver, lungs and spleen in mg normalized by body weight (in g). (C) Blood cell counts showing the concentration of erythrocytes, platelets and leukocytes. (D-E) Serum levels of alanine transaminase (ALT) (D) and aspartate transaminase (AST) (E) enzymes.

These results show that 991 encapsulation into PLGA NPs does not induce major adverse effect on blood cells and liver homeostasis.

### Chronic PLGA-991 treatment reduces inflammation in DMD muscles and improves gastrocnemius muscle homeostasis in D2-mdx mice

We measured the impact of PLGA-991 treatment over a time period of 21 days (a duration at which an improvement of muscle homeostasis and function can be observed in fibrotic DMD muscle^16^) on diaphragm and gastrocnemius muscle homeostasis (Fig.5A). First, we analyzed the inflammation level of these two muscles by flow cytometry (gating shown in Fig.S4A). The number of immune cells present within the muscle tissue was reduced by PLGA-991 treatment in both the gastrocnemius (−33.3 and -38% vs. PBS and empty PLGA, respectively) and the diaphragm (−32.6 and -23,6% *vs*. PBS and empty PLGA, respectively) (Fig.5B). More precisely, this lowered inflammation was characterized by a reduction in the number of macrophages in both muscles (−28.2 and -31.6% *vs*. PBS and empty PLGA, respectively in the gastrocnemius; -31.2 and -21.9% *vs*. PBS and empty PLGA, respectively in the diaphragm) (Fig.5C). Notably, no significant alteration of the number of neutrophils, eosinophils and T lymphocytes was observed under PLGA-991 treatment in the gastrocnemius (Fig.S4B) and the diaphragm (Fig.S4C) muscles. Next, we evaluated the consequence of PLGA-991 treatment on the dystrophic phenotype. In the gastrocnemius muscle, collagen 1 immunolabelling showed a strong reduction under PLGA-991 treatment (−20 and -15.1% *vs*. PBS and empty PLGA, respectively) (Fig.5D,F). This was associated with an increased mean cross-sectional area (CSA) of the myofibers (+29.1 and +29.6% *vs.* PBS and empty PLGA, respectively) (Fig.5E,G), leading to an improved total muscle mass (+17.3 and +14.6% *vs*. PBS and empty PLGA, respectively) Fig.5H,. However, in the diaphragm, we did not observe any improvement by PLGA-991 treatment on fibrosis (Fig.S5A), myofiber CSA (Fig.S5B) and muscle mass (Fig.S5C).

**Figure 5.**
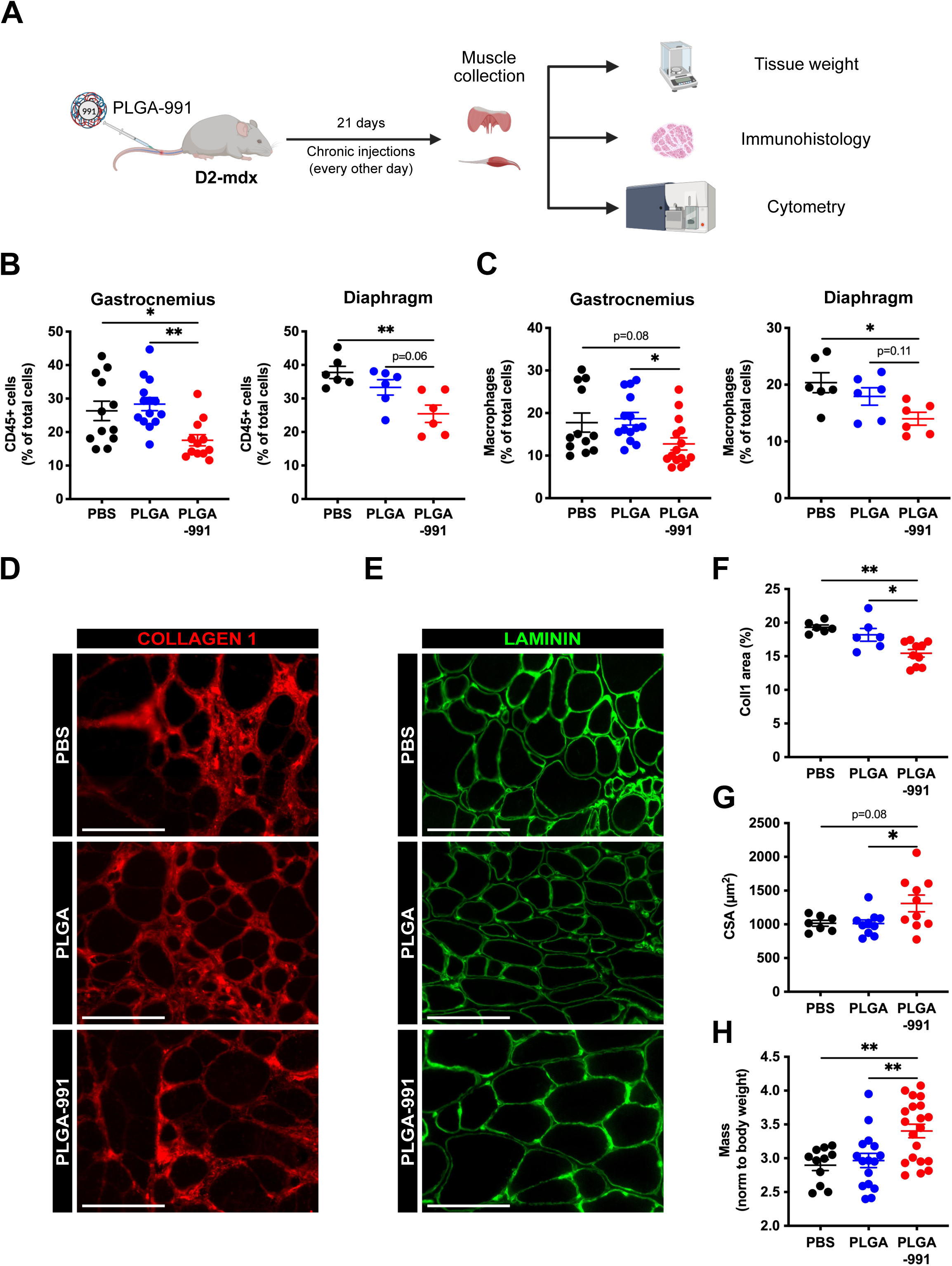
Chronic PLGA-991 treatment reduces inflammation in D2-mdx muscle and improves gastrocnemius muscle homeostasis. (A) D2-mdx mice were treated with chronic intravenous injections of PBS, PLGA or PLGA-991 for 3 weeks, then gastrocnemius and diaphragm muscles were harvested. (B,C) Muscles were digested and the proportion of immune cells (CD45+) (B) and macrophages (C) were determined by flow cytometry. (D-G) Gastrocnemius muscle sections were immunostained for Col1 (D) and Laminin (E). Scale bar: 100 μm. (F) Percentage of Col1 area. (G) Mean myofiber cross sectional area (CSA) determined on Laminin immunolabeling. (H) Relative gastrocnemius muscle mass (in mg) normalized by body weight (in g).

These results show that chronic intravenous treatment with PLGA-991 dampens muscle inflammation and improves the dystrophic phenotype in the gastrocnemius muscle of D2-mdx mice.

### Conclusions

In the present study, we established that the delivery of the AMPK activator 991 *in vivo* through its encapsulation into PLGA NPs is efficient in the D2-mdx preclinical mouse model. As compared with mdx, D2-mdx mice exhibit a more severe dystrophic phenotype characterized by fibrosis and muscle function loss in the diaphragm and gastrocnemius muscles^33–35^.

Firstly, we confirmed the useful application of microfluidic in the preparation of hydrophobic drug-loaded PLGA NPs. The translation from conventional batch mixing to microfluidic technique is a key point for achieving a more effective and easily scalable preparation method. Notably, starting from nanoprecipitation parameters, including polymer concentration, organic phase content, and aqueous/organic phase ratio, the translation into microfluidic resulted in NPs with sub-100 nm size. A DoE highlighted the crucial effects of polymer, drug concentrations and FRR on NP features. In addition, besides their optimal pharmaceutical properties (namely biocompatibility and biodegradability), their stability and the possibility to obtain ready-to-use dry powders, highlight the potential of PLGA NPs as drug delivery systems for 991.

*In vitro*, we showed that PLGA NPs were internalized by macrophages without affecting their viability, which is in accordance with previous observations^42^. Moreover, PLGA-loaded 991 activated AMPK and inhibited TGFβ secretion by fibrotic macrophages as efficiently as the free compound, suggesting that once internalized into cells, PLGA are able to release the 991 compound.

*In vivo*, biodistribution analyses in the D2-mdx mouse model indicated a high PLGA NP uptake in the liver, spleen and lungs as previously described in other models^41^. Interestingly, we also observed PLGA accumulation in the diaphragm and gastrocnemius muscles with their cellular uptake and their progressive elimination from the cells, indicative of their metabolization. The lack of fluorescence detection by imaging the whole gastrocnemius muscle, due to the low sensitivity of the method, highlights the crucial need of higher sensitivity analyses such as microscopy or flow cytometry to get a more accurate view of NP distribution. The results confirmed that previous analysis made with silica NPs^28^ and show that NPs passively enter into inflamed skeletal muscle in dystrophies, making NPs a vehicule of interest to target molecules into the diseased muscles of the whole body. A 21 days chronic injection did not impact on blood cell counts, suggesting that PLGA encapsulation of 991 compound prevents its uptake by blood cells and avoids hemolysis and platelet aggregation. Moreover, no increase in ALT and AST enzymes in blood was observed, indicative of a lack of liver injury, in accordance with the hypothesis of a low metabolization of PLGA NPs in the liver (ref?). Finally, chronic PLGA-991 treatment efficiently reduced the inflammation in both diaphragm and gastrocnemius muscles, with a specific reduction in macrophage numbers. This was associated in the gastrocnemius with decreased fibrosis and increased myofiber size and muscle mass, indicative of improved muscle homeostasis. However, despite a reduction of inflammation, no improvement of fibrosis was observed in the diaphragm. This is likely due to the more severe phenotype exhibited by the diaphragm muscle, notably the higher level of fibrosis (Fig.5E and Fig.S5A, 19.3 vs. 31.2% of Col1 area in the gastrocnemius and diaphragm muscles, respectively). Therefore, an extended treatment duration is likely required to improve diaphragm muscle homeostasis.

In conclusion, our results show that 991 encapsulation into PLGA NPs is a safe and efficient strategy for its delivery into skeletal muscle in DMD to modulate inflammation and improve muscle homeostasis.

## Materials and methods

### Reagents

PLGA 75:25 (Resomer® RG 752 H, Mw = 4–15 kDa) (analytical grade) and 378.3 g/mol D-(+)-trehalose dehydrate (trehalose) used for the freeze-drying study were purchased at Sigma-Aldrich®. 991 (molar mass, Mw = 431.87 g/mol) was ordered from Cliniscience®. Float-A-LyzerTM G2 Dialysis devices (3.5–5 kDa MWCO, 1 mL) and 1,1′-dioctadecyl-3,3,3′,3′-tetramethylindodicarbocyanine, 4-chlorobenzenesulfonate salt (DiD) were purchased from Fisher Scientific. Sodium silicotungstate used for staining in TEM was supplied by Agar Scientific). Ultra pure water was obtained using a Milli Q Academic System from Merk Millipore®. 991 (M^w^ = 431.87 g/mol) was obtained from SpiroChem AG and resuspended in methanol.

### Preparation of PLGA NPs by nanoprecipitation

Drug-loaded NPs were prepared by the nanoprecipitation technique^43^. Briefly, 6 mg of PLGA were dissolved in 800 µL of acetonitrile and added with 200 µL of a methanolic stock solution of active molecule (5 mg/mL). This organic solution was then dropped into 2 mL of Milli-Q water under magnetic stirring for 30 s. Precipitation of NPs occurred spontaneously. To remove the organic solvent and purify the NPs from the non-encapsulated drug, drug-loaded NPs were dialyzed 1 h against Milli-Q water at RT. Empty NPs, without adding drug molecule, were prepared as well. The aqueous suspension was then stored at 4 °C.

### Preparation of PLGA NPs by microfluidic technique

991-loaded PLGA NPs were obtained by microfluidic technique using benchtop NanoAssemblr^TM^ provided by Precision NanoSystem, Inc.. Disposable cartridges with a specific toroidal micromixer, with estimated channel dimensions of 500 µm (W) and 200 µm (H) were employed together with the automated NanoAssemblr^TM^. Briefly, for each preparation, to 6 mg of PLGA 75:25 dissolved in acetonitrile, an aliquot (200 µL) of a methanolic stock solution of 991 (5 mg/mL) was added in a total volume of 1 mL. The organic phase comprised of the polymer/drug solution was injected into one port of the NanoAssemblr^TM^ instrument. The aqueous phase was simultaneously injected into the second port of the system to maintain a 1:2 organic:aqueous flow rate ratio (FRR) and 12 mL·min^-1^ total flow rate (TFR). NP products were gathered in a 15 mL falcon tube, while separately disposing of the initial 0.45 mL and the final volume of 0.05 mL of NP product. To remove the organic solvent and purify the NPs from the non-incorporated drug, 991-loaded NPs were dialyzed (Spectrum^TM^ Spectra/Por^TM^ Float-A-Lyzer^TM^ G2 Dialysis devices (M^w^ = 3.5-5 kDa)) against Milli-Q water at room temperature for 1 h. Unloaded (*i.e.*, without adding 991) and DiD-loaded NPs were prepared as well. The particles were then stored at 4 °C.

### Physicochemical characterization of empty and loaded NPs

The hydrodynamic diameter, polydispersity index (PdI), and surface potential of the prepared particles were analyzed by dynamic light scattering (DLS) using a Malvern Zetasizer^®^ Nano ZS instrument (Malvern Instruments S.A., Worcestershire, UK). The angle was set at 173° and measurements were carried out at 25 °C after dilution (1:10) of the particulate suspension in Milli-Q water. The zeta potential (*ζ*) was calculated from the electrophoretic mobility measured for samples diluted in KCl 1 mM. The physical stability of NP suspensions at storage conditions (4 °C) was determined by evaluating at different interval times the mean diameter, the PdI and *ζ*. Each measurement was carried out in triplicate.

### TEM

The morphology of NPs was evaluated by TEM performed with a Philips CM120 microscope. The diluted samples (10 µL) were dropped onto a 200 mesh carbon/formvar microscope grid (copper support coated with carbon), stained with a sodium silicotungstate aqueous solution, and slowly dried in the open air. The dry samples were observed by TEM under 120 kV acceleration voltage.

### Determination of encapsulation efficiency

The encapsulation efficiency (EE) of drug-loaded PLGA NPs was determined by dissolving an aliquot of NP solution in acetonitrile to extract the encapsulated drug. The drug EE was calculated as follows:

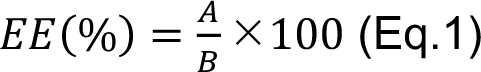

where *A* is the amount of encapsulated drug after NPs purification and *B* is the initial amount of drug determined from a fresh aliquot of organic phase. The drug loading (DL) was calculated as the ratio between the amount of entrapped drug and the total nanocarrier mass x 100. The amount of incorporated drug was determined by UHPLC. NP sample was diluted with acetonitrile (CH^3^CN)/water mixture 50/50 v/v, acidified by 0.1% trifluoroacetic acid (TFA), vortexed, and filtered through 0.22 μm Nylon filters (Agilent). The UHPLC system consisted of a quaternary pump (Waters^TM^, quaternary solvent manager-R), autosampler (Waters^TM^, sample manager FTN-R), and a multiple wavelength UV detector (Waters, 2998 PDA Detector). The data were processed using Empower 3 (Waters^TM^). The analytical column was a Cortecs C18 column (50 × 4.6 mm, 2.7 μm; Waters) at 40 °C; the mobile phase consisted of waters 0.1% TFA (solvent A) and CH^3^CN 0.1% TFA (solvent B) at a flow rate of 0.5 mL/min with isocratic conditions: 40% A and 60% B. The drug concentration in the column effluent was monitored by its absorbance at 254 nm. The quantification of the active molecule was done using calibration with standard solutions chromatographed under the same experimental conditions, with a concentration range of 0.25–200 μg/mL. The calibration curve was linear (*R*^2^ > 0.995).

### NP optimization using a design of experiment (DoE)

A 2^3^ full factorial design was used to investigate the effect of 3 experimental factors (PLGA, drug concentrations and *FRR*) and their interactions on NP physicochemical characteristics. For each factor, the experimental domain whose lower and upper limits correspond to the coded levels −1 and +1, respectively was defined. The center point corresponds to the coded level 0. As reported in Table 1, the polymer concentration (*X*^1^) varied from 5–10 mg/mL, *FRR* (*X*^2^) from 1:2–1:5 and drug amount (*X*^3^) from 0.7–1.4 mg/mL. The studied responses were the encapsulation efficiency (*Y*^1^) and the drug loading (*Y*^2^) as NP characteristics to be well-controlled.

**Table 1.** Selected experimental factors and their coded levels in the 2^3^ full factorial design

| Experimental factors | Coded levels |  |  |
| --- | --- | --- | --- |
|  | - 1 | 0 | + 1 |
| PLGA conc. (mg/mL)<br>($X_1$) | 5 | 7.5 | 10 |
| *FRR* ($X_2$) | 1:5 | 1:3.5 | 1:2 |
| Drug conc. (mg/mL)<br>($X_3$) | 0.7 | 1.1 | 1.4 |

The full factorial design consisted in 2^3^ runs defined as every possible combination of +1 and −1 levels selected in turn and allowed calculating a synergistic model, as follows:

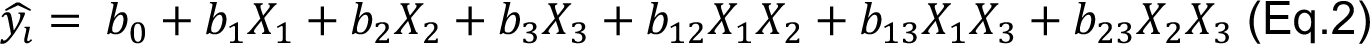

where 𝑦̂_*i*_ is the predicted response for experiment i, *b*^0^ is the intercept, the coefficients *b*^ì^ correspond to the main effects of the experimental factors (*X*^i^) and the coefficients *b*^ij^ are the effects of the two-factor interactions.

Two center points were also added in order to estimate the variance of the experimental error and to evaluate the predictive performance of the developed models. The corresponding experimentation plan is presented in Table S2. Multiple linear regression calculations, statistical analysis and response contour plots were performed with Umetrics MODDE 12.0 software.

### Freeze-drying of drug-loaded NPs

Drug-loaded NPs were freeze-dried using the CRYONEXT pilot freeze dryer (Cryotec). The suspension was prepared by diluting the formulation 1/5 with a solution of trehalose at a final concentration of 12% w/v. The freeze-drying process started reaching an initial cool temperature of −35 °C for 3 h. Then, the temperature of the shelf was increased to -20 °C for 10 h with a chamber pressure of 0.25 mBar to remove ice by sublimation (primary drying). Secondary drying was carried out for 5 h at 4 °C, with 0.1 mBar pressure. Finally, freeze-dried NPs were resuspended in 1 mL of Milli-Q water and vortexed until complete dissolution of the pellet. Size, PdI, ζ, EE, morphology and storage physical stability at 4 °C of NPs were measured after resuspension. The osmolarity of 50 µL of reconstituted sample was measured at RT using a Gonotec Osmomat 3000 provided by Thermo Fisher Scientific^®^ (Waltham, USA).

### Determination and quantification of organic solvent residuals

NPs were dialyzed for 1 h against Milli-Q water. The quantifications of remaining acetonitrile were performed by ^1^H NMR analysis. ^1^H NMR spectra were recorded on the liquid samples in 5 mm tubes with a liquid-state high-resolution Bruker AVL300 NMR spectrometer operating at 300 MHz under quantitative analysis conditions. The FID was recorded during an acquisition time of 5.5 s after a 90° radiofrequency pulse (14 µs). A delay of 60 s was applied between each scan for ensuring full relaxation of the FID. 8 scans were accumulated with conventional phase cycling. 500 µL of an aqueous suspension of NPs were added to 500 µL of deuterium oxide (D^2^O) containing 2% v/v of acetone used as internal standard. Quantification of acetonitrile residuals was done from the integral of the acetonitrile peak at a chemical shift of 1.98 ppm and that of the acetone reference peak at 2.15 ppm. The quantification of acetonitrile was performed on lyophilized samples as well.

### Mice experiments

D2-mdx (D2.B10-Dmd^mdx^/J) mice were bred and used according to French legislation, the protocols were approved by the local Ethical committee and Ministery from. Experiments were conducted on males of 8-16 weeks of age.

### Murine BMDM isolation and culture

Macrophages were derived from murine bone marrow precursors as previously described^18^and were cultured in DMEM containing 20% heat inactivated Fetal Bovine Serum (FBS), 30% of L929 cell line-derived conditioned medium (enriched in CSF-1), 2.5 μg/mL of fungizone and 100 U/mL of penicillin/streptomycin for 6-7 days. Macrophages were then activated into fibrotic cells with 1 µg/mL of total protein lysate isolated from tibialis anterior of fibrotic mdx (Fib-mdx) mouse model^44^, in DMEM containing 10% FBS for 2 days, as previously described^16^. For viability assessment, BMDMs were labeled with Alexa Fluor 488-conjugated Annexin V (A13201, Molecular Probes) and DAPI (564907, BD Biosciences) and analyzed on a BD FACSCanto II apparatus.

### NP internalization by BMDMs

BMDMs were seeded at 54,000 cells/cm^2^ in DMEM containing 10% of FBS on a coverslip in 12-well plates. Activated fibrotic macrophages were incubated with DiD-loaded PLGA NPs at the final DiD concentration of 0.9 µg/mL for 0, 2, 4 and 24 h at 37 °C. After incubation, cells were fixed for 10 min in 4% formaldehyde, permeabilized 10 min in 0.5 % Triton X-100 and saturated in 2% BSA for 1 h at room temperature. They were then labeled with anti-F4/80 primary antibody (E-AB-F0995A, Elabscience, 1:200) overnight at 4 °C followed by an incubation with a secondary antibody coupled with Cy3 (Jacson Immunoresearch, 1:200) for 2 h at 37 °C. After nuclei staining with Hoechst 33342 (Sigma), coverslips were mounted in Fluoromount (Interchim) and imaged on a TCS SP5-X confocal microscope (Leica microsystems). Images were analyzed with Las X software (Leica Microsystems)

### TGFβ1 quantification

BMDMs were seeded at 100,000 cells/cm^2^ in 48-well plates in DMEM containing 10% of FBS. Activated fibrotic macrophages were incubated for 20h with 10 µM 991, empty PLGA NPs or PLGA-991. Vehicle control (Veh) consisted of a treatment with an equivalent concentration of methanol. After 2 washes with PBS, were cultured for 20 h in DMEM containing 0.1% BSA. The conditioned medium was then harvested and centrifuged for 5 min at 500 g to remove cell debris. The total amount of TGFβ1 was determined on technical duplicates by ELISA (88-8350-22, Invitrogen), after heat denaturation followed by acidic treatment and neutralization according to the manufacturer’s instructions.

### Western blot

BMDMs were seeded in DMEM containing 10% of FBS at 208,000 cells/cm^2^ in 6-well plates and activated into fibrotic macrophages. One hour before treatment, culture medium was replaced by DMEM without red phenol (Gibco) supplemented with 10% charcoal stripped FBS (Gibco) and cells were treated with 10 or 20 µM 991, empty PLGA NPs or PLGA-991 for 6 h. Proteins were isolated in lysis buffer containing 50 mM Tris-HCL (pH 7.5), 1 mM EDTA, 1 mM EGTA, 0.27 M sucrose, 1% triton, 20 mM glycerol-2-phosphate disodium, 50 mM NaF, 0.5 mM PMSF, 1 mM benzamidine, 1 mM Na3VO4 and 1% cocktail phosphatase inhibitor 3 (Sigma-Aldrich, P0044) for 30 min on ice and centrifuged for 10 min at 16,100 g to remove debris. Ten mg of proteins were subjected to SDS-PAGE and transferred onto a nitrocellulose membrane which was probed with antibodies against p-ACC (#3661, Cell Signaling, 1:1000), ACC (#3676, Cell Signaling, 1:1000), p-RAPTOR (#, 1:1000), RAPTOR (#, 1:1000) or b-actin (#A5316, Sigma-Aldrich, 1:5000). Blots were revealed on a Chemidoc imager (Bio-Rad) using SuperSignal West Femto Maximum Sensitivity Substrate (Thermo Scientific) and signal intensity was quantified using Image Lab Software (Bio-Rad).

### Organ biodistribution

200 µL of DiD-loaded NPs were administered to anesthetized mice by intravenous injection in the tail vein. At scheduled time points (0.5, 1, 24, 48, 72 h), anesthetized animals were fully imaged in a light-tight chamber where a controlled flow of 1.5% isoflurane in the air was administered to maintain anesthesia. Fluorescence images, as well as bright-field pictures of mice’s whole body (ventral and prone view), were acquired *via* a back-thinned CCD-cooled camera ORCA II-BT-512G (Hamamatsu Photonics) using a coloured glass long-pass RG 665 filter (Melles Griot), which cuts off all excitation light. Optical excitation was carried out at 644 nm, and the emission wavelength was detected at 655 nm. At each time point, mice were sacrificed and tissues and organs (lungs, liver, spleen, diaphragm and gastrocnemius) were harvested for *ex vivo* imaging. Images were analyzed using the Wasabi software 1.5 (Hamamatsu Photonics).

### Chronic PLGA-991 intravenous injection

200 µL empty PLGA NPs or PLGA-991 were administered every other day to anesthetized mice by intravenous injection in the tail vein (3.5 mg/kg of drug). Control mice were injected similarly with the same volume of PBS.

### Blood cell count

Blood was collected by cardiac puncture directly after sacrifice and transferred into a tube previously coated with PBS containing 50 mM EDTA. Blood was diluted in PBS and erythrocytes were counted on a hemocytometer. For platelet numeration, blood was diluted in 1% ammonium oxalate (Sigma) and incubated on a hemocytometer for 15 min in a humid chamber before counting. Leukocytes were counted on a hemocytometer after blood was diluted in Türk’s solution (Sigma).

### Plasma transaminase quantification

Blood was centrifugated at 1,500 g for 15 min and plasma was aliquoted and stored at -80°C for further analysis. Alanine transaminase (CAK1002, Cohesion Biosciences) and Aspartate transaminase (CAK1004, Cohesion Biosciences) levels were determined on technical duplicates of respectively 10 and 20 µL of plasma according to manufacturer’s instructions.

### Muscle flow cytometry analysis

Muscles were dissociated and digested in DMEM F/12 medium (GIBCO) containing 10 mg/ml of collagenase B and 2.4 U/mL Dispase II (Roche Diagnostics GmBH) at 37°C for 1 h, passed through a 30 µm cell strainer and erythrocytes were lysed using ACK lysis buffer (Lonza). For analysis of DiD-loaded NPs internalization, cells were labeled with DAPI (564907, BD Biosciences) and analyzed on a BD FACSCanto II (BD Biosciences) apparatus. For immune cells characterization, dead cells were first labeled with Ghost Dye Red 780 (13-0865-T500, Tonbo Biosciences) for 30 min at 4°C, followed by incubation with anti-mouse FcR blocking reagent (130-092-575, Miltenyi Biotec) in PBS 2% FBS for 10 min at 4°C. Cells were then stained with BV510-conjugated anti-CD45 (103138, Biolegend), PE-Cy7-conjugated anti-CD64 (139314, Biolegend), APC-conjugated anti-Ly-6G (17-9668, eBioscience), FITC-conjugated anti-CD3 (11-0032-82, eBioscience) and PE-conjugated anti-Siglec-F (552126, BD Biosciences) antibodies for 30 min at 4°C and analyzed on a BD FACSCanto II apparatus.

### Histology and immunofluorescence analyses in mouse

Diaphragm and gastrocnemius muscles were frozen in nitrogen-chilled isopentane and kept at -80°C until use. Ten µm-thick cryosections were prepared, permeabilized 10 min in 0.5 % Triton X-100 and saturated in 2% bovine serum albumin (BSA) for 1 h at room temperature. Sections were incubated with primary rabbit anti-Laminin (L9393, Sigma Aldrich, 1:200) and goat anti-Collagen I (131001, Biotech, 1:400) antibodies overnight at 4°C, then labeled with secondary donkey anti-rabbit and anti-goat antibodies were coupled to FITC and Cy3, respectively (Jackson Immunoresearch, 1:200). Fluorescent immunolabelings were recorded with a Zeiss Axio Scan.Z1 microscope connected to an ORCA-Flash4.0 V2 CMOS camera (Hamamatsu Photonics) at 20X magnification. Areas of Collagen1 were calculated with ImageJ software as previously described^16^ on 8-10 fields randomly chosen. Cross Section Area (CSA) was determined on whole muscle sections using Open-CSAM program as previously described^45^.

### Statistical analyses

All experiments were performed using at least 3 independent different cultures or animals except for Fig.2E where two mice were used. Results are shown as mean+/-sem. Depending on the experimental, one-way or two-way ANOVA with multiple comparisons were performed with GraphPad Prism version 9.0.0 for statistical analyses. *p<0.05, **p<0.01, ***p<0.001.

## Acknowledgements

This work was funded by Agence Nationale de la Recherche (ANR-19-CE14-0008 and ANR-18-CE18-0025-01). GJ and AK were supported by AFM-Telethon (MyoNeurALP Alliance).

## Author contribution

Conceptualization: DK, BS, SA, CB, YC, BC, RM, GL, GJ ; Methodology: IA, AK, SBL, FT, AF, JSB, GJ ; Analysis: IA, AK, SBL, FT, AF, JSB, GJ ; Writing original draft: IA, GJ; Writing review & editing: IA, AK, DK, BS, SA, CB, YC, BC, RM, GL, GJ ; Supervision: BC, RM, GL, GJ ; Funding Acquisition: BC, RM, GL, GJ.

## Figure legends

**Supplementary figure 1.**
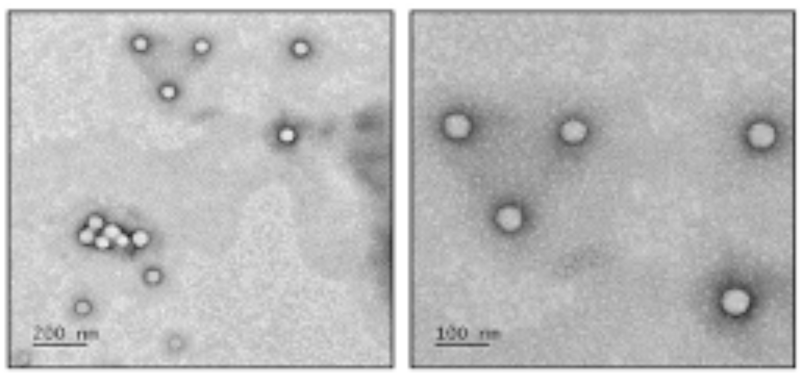
Effect of freeze-drying on PLGA-991 NP morphology. TEM images of freeze-dried 991-loaded PLGA NPs prepared by microfluidic technique after resuspension.

**Supplementary figure 2.**
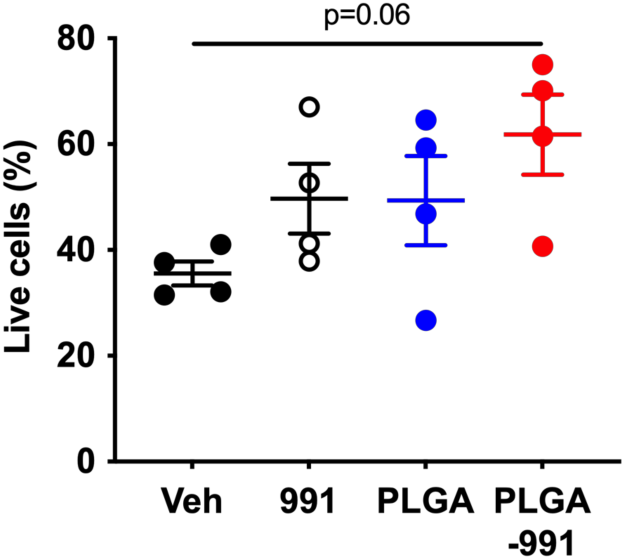
Effect of PLGA-991 on macrophage viability *in vitro*. BMDMs were polarized into fibrotic macrophages and treated with 20 µM of 991 alone, empty PLGA or PLGA-991 for 20 h and cell viability was assessed by flow cytometry after Annexin V and Hoechst labeling. The percentage of viable cells, identified as Annexin V^neg^ Hoechst^neg^, is shown.

**Supplementary figure 3.**
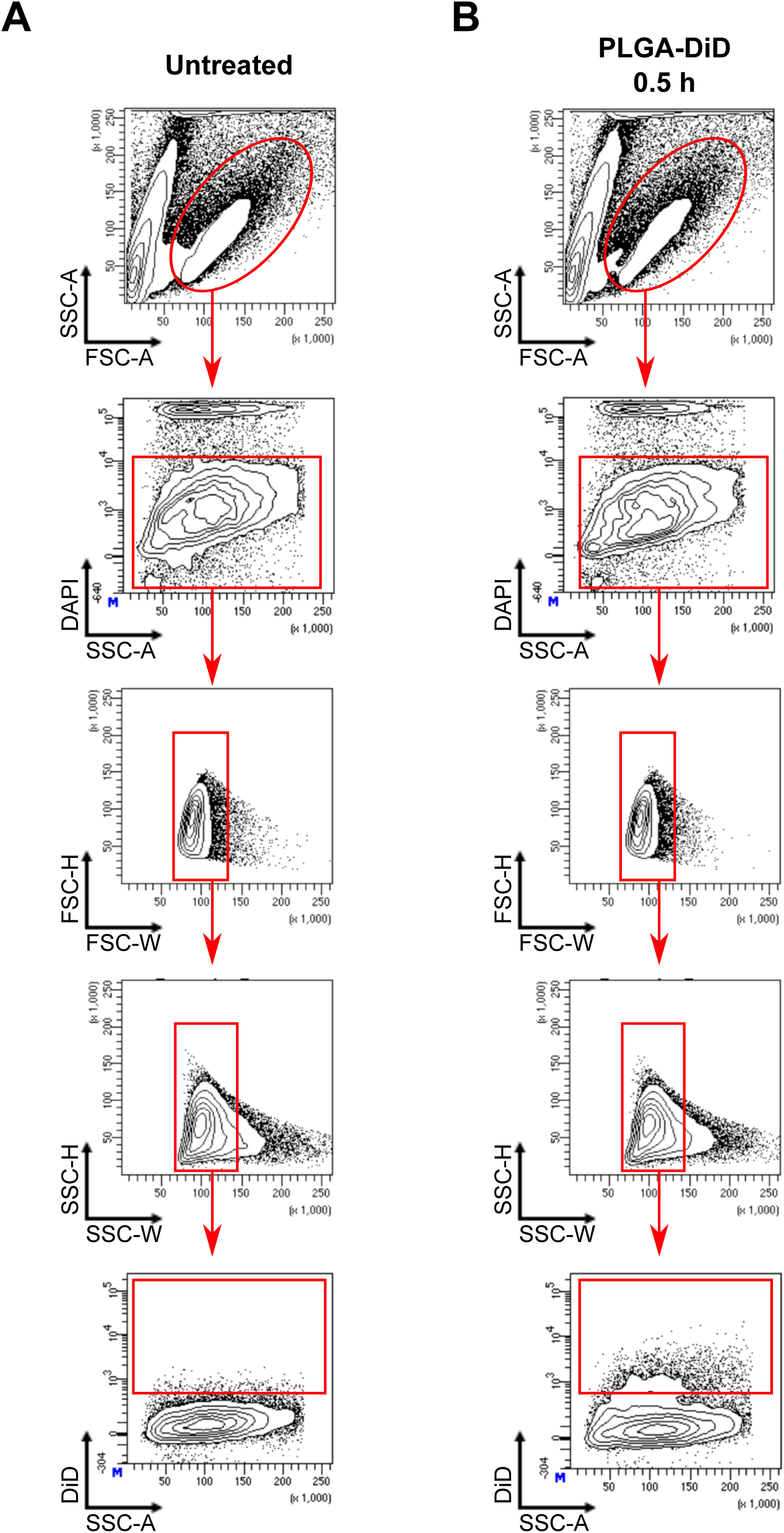
Gating strategy to quantify NP internalization in muscle cells. D2-mdx mice were injected intravenously with DiD-loaded PLGA NPs as in Fig.3A. Muscles were digested using Collagenase B and cells were labeled with DAPI. Representative plots obtained on the diaphragm muscle at 0.5h post-injection are shown.

**Supplementary figure 4.**
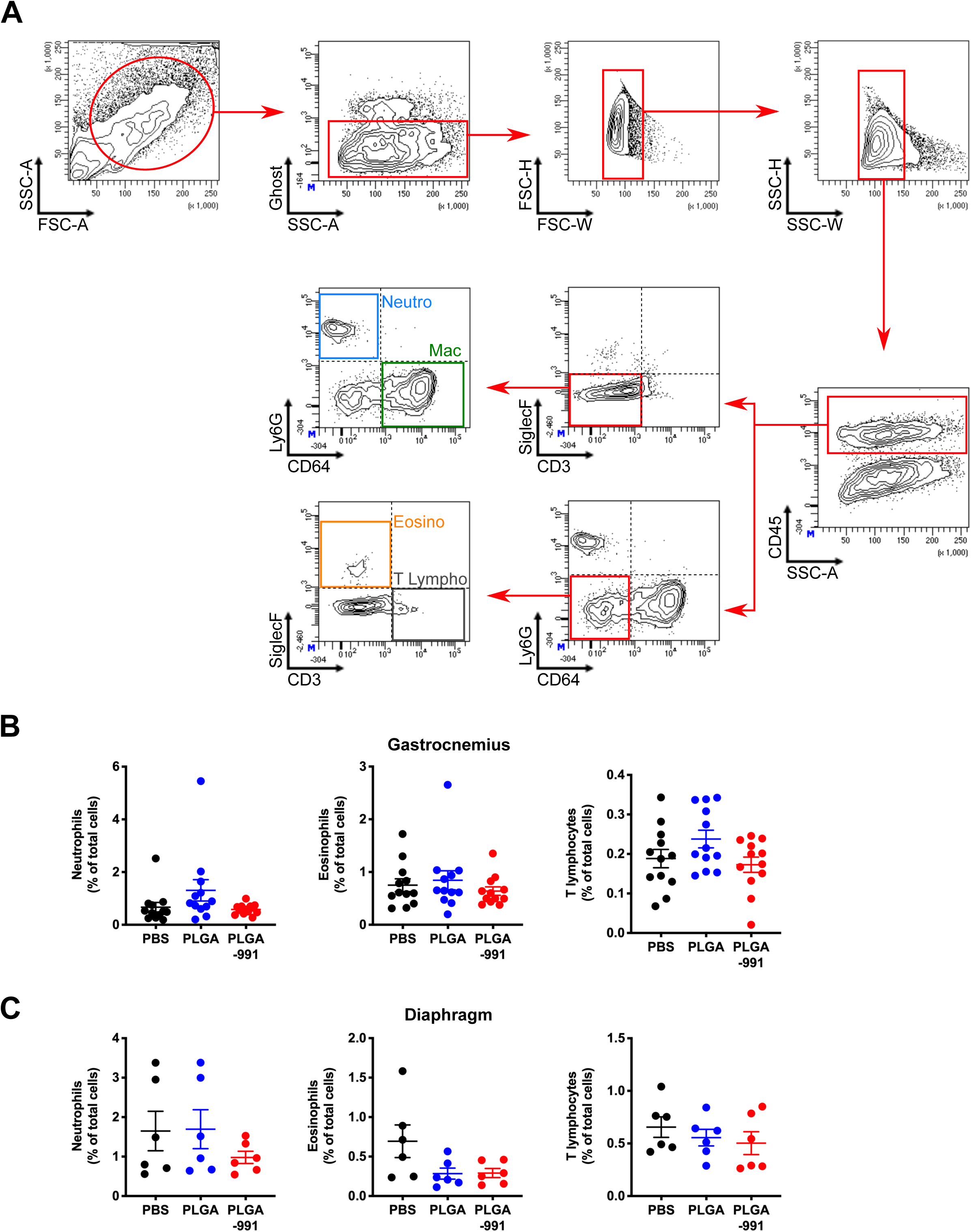
Immune cell characterization in the muscles. D2-mdx mice were treated with chronic intravenous injections of PBS, PLGA or PLGA-991 for 21 days as in Fig.5A, then gastrocnemius and diaphragm were collected and were digested using Collagenase B. Cells were blocked with anti-mouse FcγRII/III and labeled with CD45, CD64, Ly6G, CD3 and SiglecF antibodies to identify macrophages (Mac; CD45+CD64+Ly6G-CD3-SiglecF-), neutrophils (Neutro; CD45+CD64-Ly6G+CD3-SiglecF-), eosinophils (Eosino; CD45+CD64-Ly6G-CD3-SiglecF+) and T lymphocytes (T Lympho; CD45+CD64-Ly6G-CD3+SiglecF-) by flow cytometry. Ghost 780 dye was used to exclude dead cells. (A) Representative plots obtained on gastrocnemius muscle from PBS-treated mice. (B-C) Quantification of neutrophils, eosinophils and T lymphocytes in the gastrocnemius (B) and diaphragm (C) muscles.

**Supplementary figure 5.**
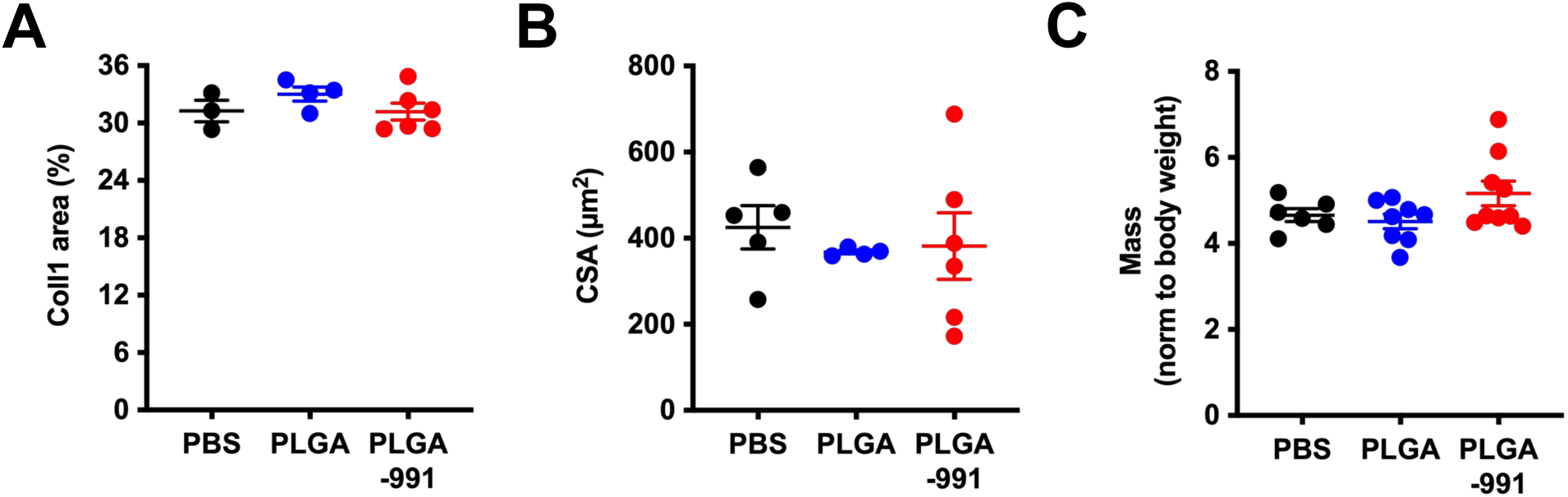
Effect of PLGA-991 chronic administration on diaphragm muscle homeostasis. D2-mdx mice were treated with chronic intravenous injections of PBS, PLGA or PLGA-991 for 21 days as in Fig.5A, and diaphragm was collected. (A) Percentage of fibrotic areas determined on muscle sections immunostained for Col1. (B) Mean CSA of myofibers determined on whole muscle sections after laminin immunolabeling. (C) Relative gastrocnemius muscle mass (in mg) normalized by body weight (in g).

**Supplementary Table 1.**
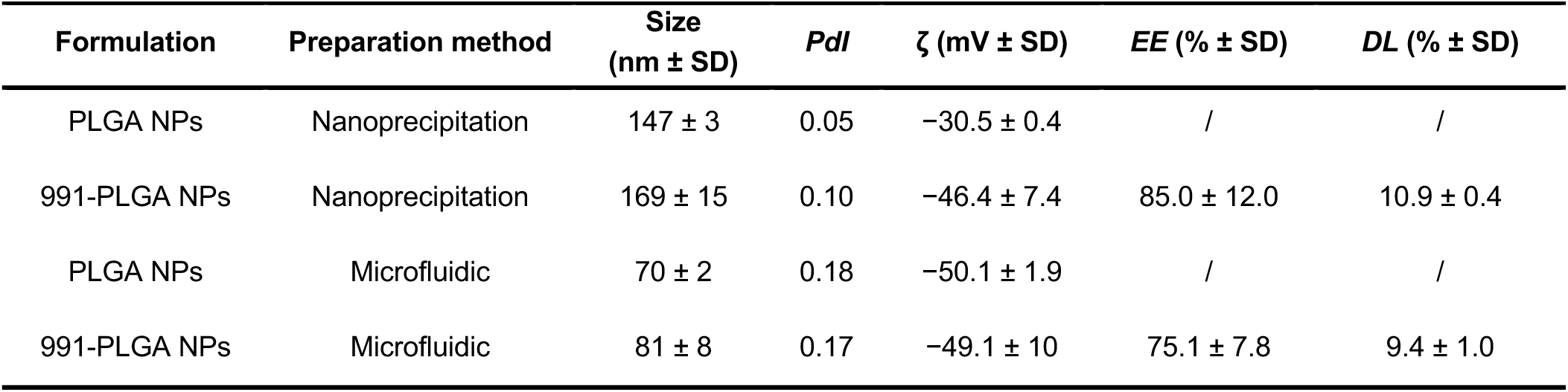
Physicochemical characterization of blank and loaded PLGA NPs prepared by nanoprecipitation and microfluidic technique. Values are given as mean ± SD (n = 3). NPs nanoparticles; PdI: polydispersity index

| Formulation | Preparation method | Size<br>(nm ± SD) | *Pdl* | $\zeta$ (mV ± SD) | *EE* (% ± SD) | *DL* (% ± SD) |
| --- | --- | --- | --- | --- | --- | --- |
| PLGA NPs | Nanoprecipitation | 147 ± 3 | 0.05 | -30.5 ± 0.4 | / | / |
| 991-PLGA NPs | Nanoprecipitation | 169 ± 15 | 0.10 | -46.4 ± 7.4 | 85.0 ± 12.0 | 10.9 ± 0.4 |
| PLGA NPs | Microfluidic | 70 ± 2 | 0.18 | -50.1 ± 1.9 | / | / |
| 991-PLGA NPs | Microfluidic | 81 ± 8 | 0.17 | -49.1 ± 10 | 75.1 ± 7.8 | 9.4 ± 1.0 |

**Supplementary Table 2.**
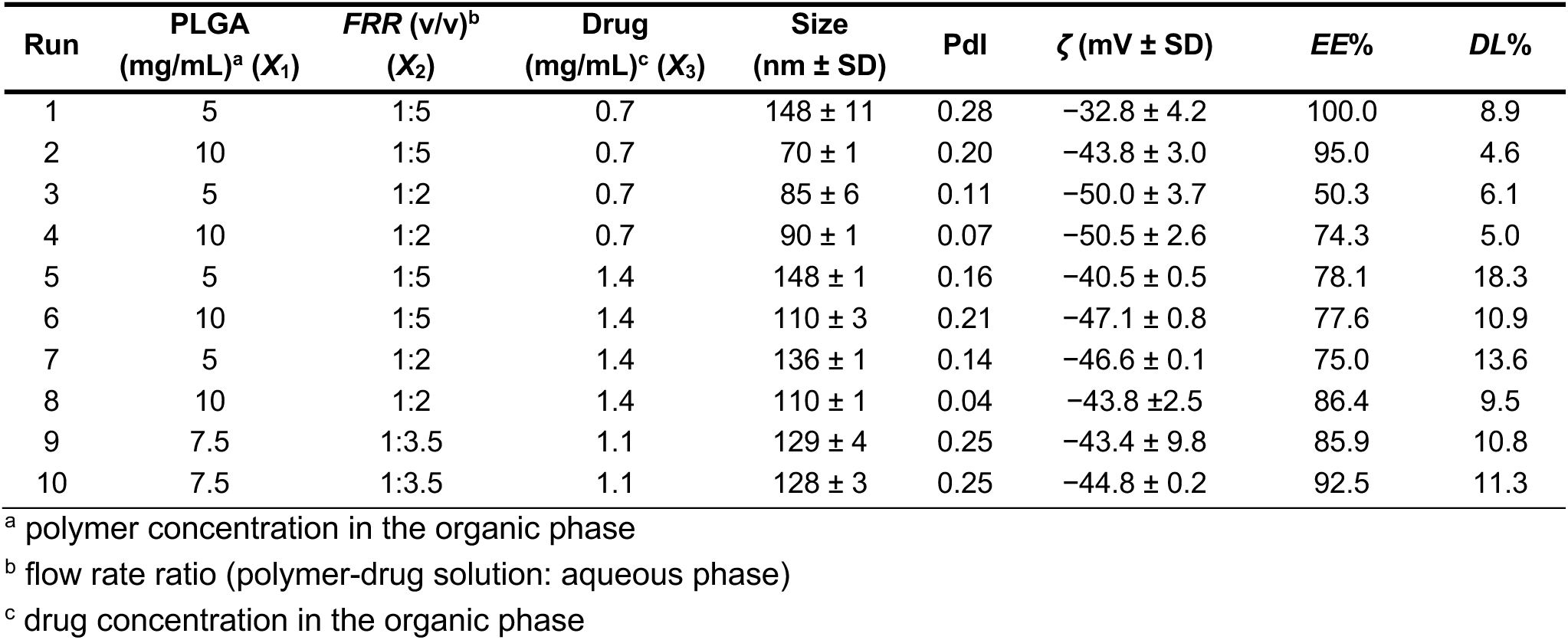
Experimentation plan built according to a 2^3^ full factorial design (runs 1 to 8) with two additional center points (runs 9 and 10) with the corresponding responses EE and DL.

**Supplementary Table 3.**
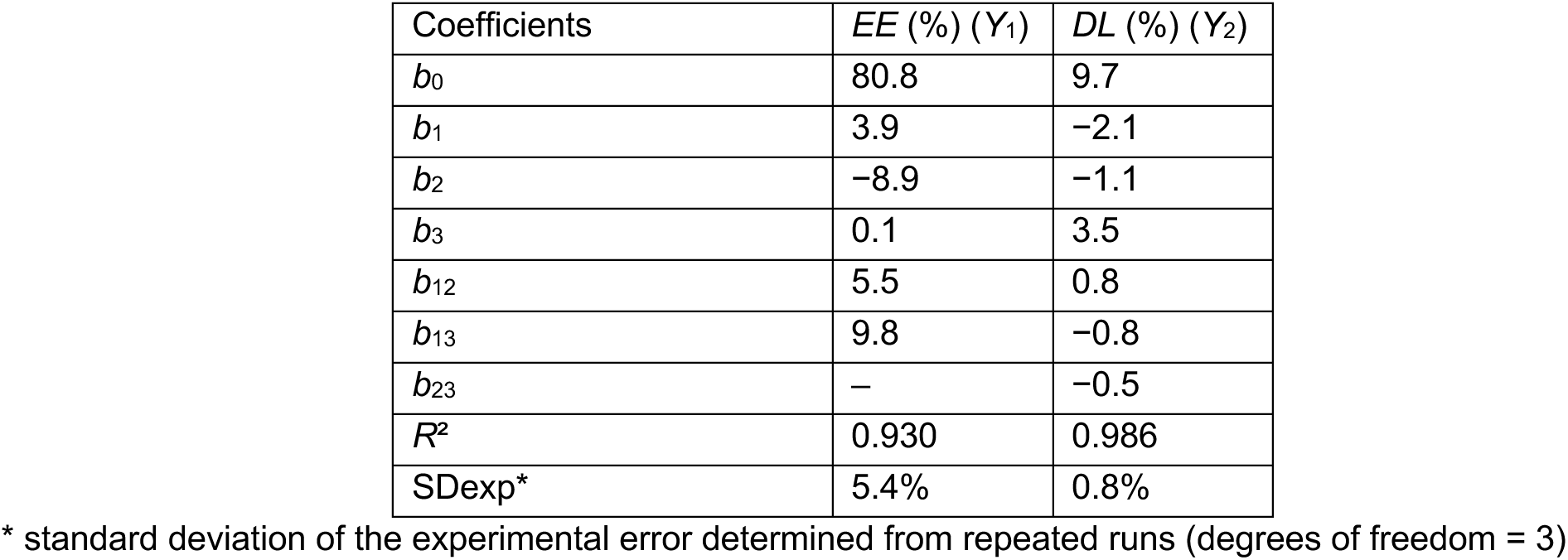
Coefficients of synergistic models (Eq. 2) for EE and DL with their corresponding R^2^ and experimental standard deviation SDexp.

**Supplementary Table 4.**
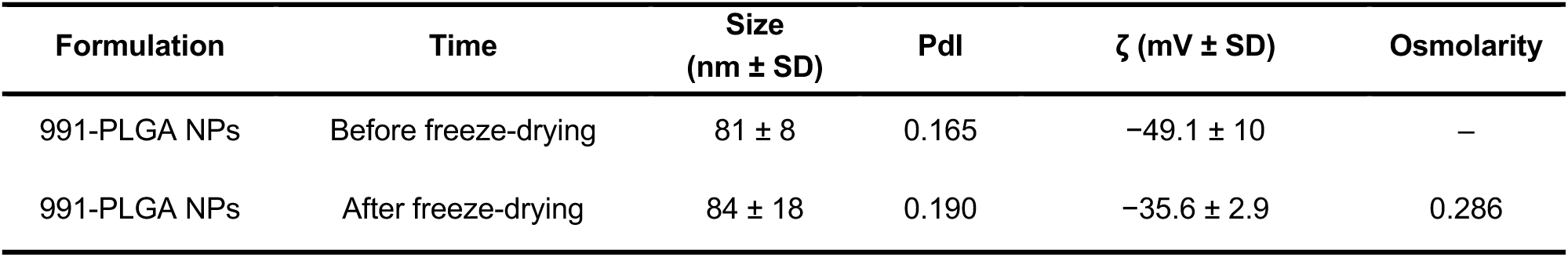
Physicochemical characterization of loaded-PLGA NPs prepared by microfluidic technique, before and after freeze-drying (n=3).

