## Supplementary tables for "Polymeric nanoparticles delivery of AMPK activator 991 prevents its toxicity and improves muscle homeostasis in Duchenne Muscular Dystrophy"

**Supplementary Table 1**: Physicochemical characterization of blank and loaded PLGA NPs prepared by nanoprecipitation and microfluidic technique. Values are given as mean ± SD (n = 3). NPs: nanoparticles; PdI: polydispersity index; EE: Encapsulation Efficiency; DL: Drug Loading.

| **Formulation** | **Preparation method** | **Size (nm ± SD)** | ***PdI*** | **ζ (mV ± SD)** | ***EE* (% ± SD)** | ***DL* (% ± SD)** |
|---|---|---|---|---|---|---|
| PLGA NPs | Nanoprecipitation | 147 ± 3 | 0.05 | −30.5 ± 0.4 | / | / |
| 991-PLGA NPs | Nanoprecipitation | 169 ± 15 | 0.10 | −46.4 ± 7.4 | 85.0 ± 12.0 | 10.9 ± 0.4 |
| PLGA NPs | Microfluidic | 70 ± 2 | 0.18 | −50.1 ± 1.9 | / | / |
| 991-PLGA NPs | Microfluidic | 81 ± 8 | 0.17 | −49.1 ± 10 | 75.1 ± 7.8 | 9.4 ± 1.0 |

**Supplementary Table 2.** Experimentation plan built according to a $2^3$ full factorial design (runs 1 to 8) with two additional center points (runs 9 and 10) with the corresponding EE and DL responses (see legends in Table S2).

| Run | PLGA (mg/mL)[a] ($X_1$) | *FRR* (v/v)[b] ($X_2$) | Drug (mg/mL)[c] ($X_3$) | Size (nm ± SD) | PdI | ζ (mV ± SD) | *EE*% | *DL*% |
|---|---|---|---|---|---|---|---|---|
| 1 | 5 | 1:5 | 0.7 | 148 ± 11 | 0.28 | −32.8 ± 4.2 | 100.0 | 8.9 |
| 2 | 10 | 1:5 | 0.7 | 70 ± 1 | 0.20 | −43.8 ± 3.0 | 95.0 | 4.6 |
| 3 | 5 | 1:2 | 0.7 | 85 ± 6 | 0.11 | −50.0 ± 3.7 | 50.3 | 6.1 |
| 4 | 10 | 1:2 | 0.7 | 90 ± 1 | 0.07 | −50.5 ± 2.6 | 74.3 | 5.0 |
| 5 | 5 | 1:5 | 1.4 | 148 ± 1 | 0.16 | −40.5 ± 0.5 | 78.1 | 18.3 |
| 6 | 10 | 1:5 | 1.4 | 110 ± 3 | 0.21 | −47.1 ± 0.8 | 77.6 | 10.9 |
| 7 | 5 | 1:2 | 1.4 | 136 ± 1 | 0.14 | −46.6 ± 0.1 | 75.0 | 13.6 |
| 8 | 10 | 1:2 | 1.4 | 110 ± 1 | 0.04 | −43.8 ±2.5 | 86.4 | 9.5 |
| 9 | 7.5 | 1:3.5 | 1.1 | 129 ± 4 | 0.25 | −43.4 ± 9.8 | 85.9 | 10.8 |
| 10 | 7.5 | 1:3.5 | 1.1 | 128 ± 3 | 0.25 | −44.8 ± 0.2 | 92.5 | 11.3 |

[a] polymer concentration in the organic phase

[b] flow rate ratio (polymer-drug solution: aqueous phase)

[c] drug concentration in the organic phase

**Supplementary Table 3.** Coefficients of synergistic models (Eq. 2) for EE and DL with their corresponding $R^2$ and experimental standard deviation $SD_{exp}$.

| Coefficients | *EE* (%) ($Y_1$) | *DL* (%) ($Y_2$) |
|---|---|---|
| $b_0$ | 80.8 | 9.7 |
| $b_1$ | 3.9 | −2.1 |
| $b_2$ | −8.9 | −1.1 |
| $b_3$ | 0.1 | 3.5 |
| $b_{12}$ | 5.5 | 0.8 |
| $b_{13}$ | 9.8 | −0.8 |
| $b_{23}$ | – | −0.5 |
| $R^2$ | 0.930 | 0.986 |
| $SD_{exp}$* | 5.4% | 0.8% |

* standard deviation of the experimental error determined from repeated runs (degrees of freedom = 3)

**Supplementary Table 4.** Physicochemical characterization of loaded-PLGA NPs prepared by microfluidic technique, before and after freeze-drying (n=3).

| Formulation | Time | Size (nm ± SD) | PdI | ζ (mV ± SD) | Osmolarity |
|---|---|---|---|---|---|
| 991-PLGA NPs | Before freeze-drying | 81 ± 8 | 0.165 | −49.1 ± 10 | – |
| 991-PLGA NPs | After freeze-drying | 84 ± 18 | 0.190 | −35.6 ± 2.9 | 0.286 |
